# Women’s representation in Indian academia and conferences

**DOI:** 10.1101/2023.10.26.564078

**Authors:** Shruti Muralidhar, Vaishnavi Ananthanarayanan

## Abstract

Indian science academia has a dearth of women researchers at all levels. Not only are they under-represented, but they are also under-highlighted, under-mentored and overlooked for awards, grants and other career-advancing steps. To effectively address this problem and devise a solution for the inequity, we need data on the proportion of women faculty across multiple STEM institutions. Such a database, currently, does not exist. To fill this gap, we formed BiasWatchIndia to (1) document the inequities, and (2) provide real-time actionable data as a basis for future remedial steps. Along with collecting data on women representation at the faculty level in Indian STEM (science, technology, engineering and mathematics) academia, BiasWatchIndia also helps highlight the lack of women representation and gender imbalance in Indian STEM talks, conferences, workshops and panels. Based on our findings, we recommend several measures that need to be implemented by universities and institutes to challenge the *status quo* changes for women in Indian academia.

## Methods

The proportion of women faculty across 100 universities and institutes were manually collected and curated by volunteers and interns. Each volunteer/intern visited the website of the institute or university and navigated to each of the department pages which typically hold the list of all the faculty. The total number of faculty members was counted and the fraction of women faculty was also noted. For each woman faculty, career stage details were also noted. We calculated the proportion of women faculty members at each career stage by dividing the number of women faculty members at a given stage by the total number of faculty members at that stage. All faculty were classified into the following categories based on their research areas - Biology, Physics, Chemistry, Mathematics, Engineering, Computer Sciences and Earth Sciences. All Bio-related department faculty members were put under Biology (Ecology, Cell Biology, Biophysics, etc.). Similarly, Chemistry included Organic, Inorganic, Materials Research. Physics included High Energy Physics, Physics, etc. All Engineering allied departments (Chemical, Mechanical, Electrical and Electronics, Textile, etc.) were clubbed together, including the Design department. Computer Science or a Computer Science and Engineering department were combined under Computer Science, as were AI and Machine learning, Computational Data Systems. Earth Sciences included departments such as Renewable Technology, Earth Sciences, Water Research, etc.

The proportion of women speakers for a conference or meeting was calculated based on the poster announcements that typically contain the speaker list.

Base rate data were collected between June 2020 and Dec 2021, Conference data were collected and analyzed in two stages - between June 2020 and Aug 2021 followed by Aug 2021 to March 2023.

## Results

### 1. Base rates of women faculty members in Indian academia

We first sought to estimate the proportion of women across different STEM disciplines in India. We define this ratio of number of women faculty members to the total number of faculty members as the ‘base rate’. Since these estimates were not available publicly, we calculated the base rates of women faculty members (see Methods) for seven different fields, namely Biology, Mathematics, Earth Sciences, Physics, Computer Science, Chemistry and Engineering across. We surveyed 98 universities and institutes across the country and observed that the total base rate across all the fields had a median of 16.7% (*or 0*.*17*) (**Fig. 1**). The other fields had median base rates ranging from Biology with the highest of 22.5% (*0*.*23*) to Engineering with the lowest at 8.3% (*0*.*08*) (**Fig. 1**).

**Figure 1.**
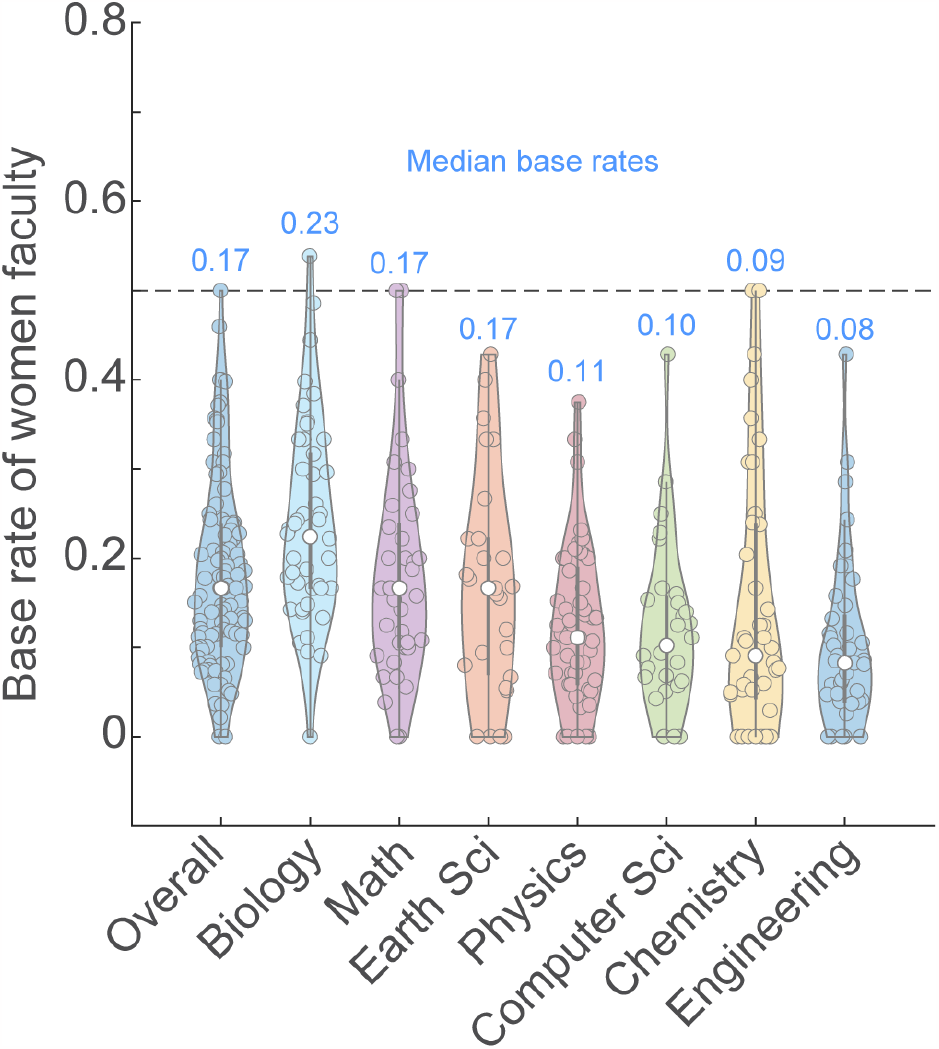
The overall base rate of women faculty in Indian academia is 16.6%. Violin plot of the base rates of women faculty members in Indian academia by field. The median base rate is indicated by the white circle and mentioned above the specific field. The thick gray lines represent the interquartile ranges. For calculation of the overall base rate, n=98 universities and institutes were surveyed, and for the field-specific base rate calculations, n=46,37,27,49,28,41 and 41 respectively for Biology, Math, Earth Sciences, Physics, Computer Science, Chemistry and Engineering.

Previous reports from the UK have noted a negative correlation between the ranking of a university and the proportion of women and members of marginalized communities hired by these universities^1^, and a higher gender disparity in pay^2^. We therefore tested the former in our dataset by plotting the base rate of the top 8 institutes (which were part of our base rate dataset) according to the National Institutional Research Framework (NIRF) rankings 2022^3^ (**Fig. 2**). Indeed, all 8 of these institutes had women base rates less than the median overall base rate of 16.6%. The median base rate of these top institutes/universities was only 10%.

**Figure 2.**
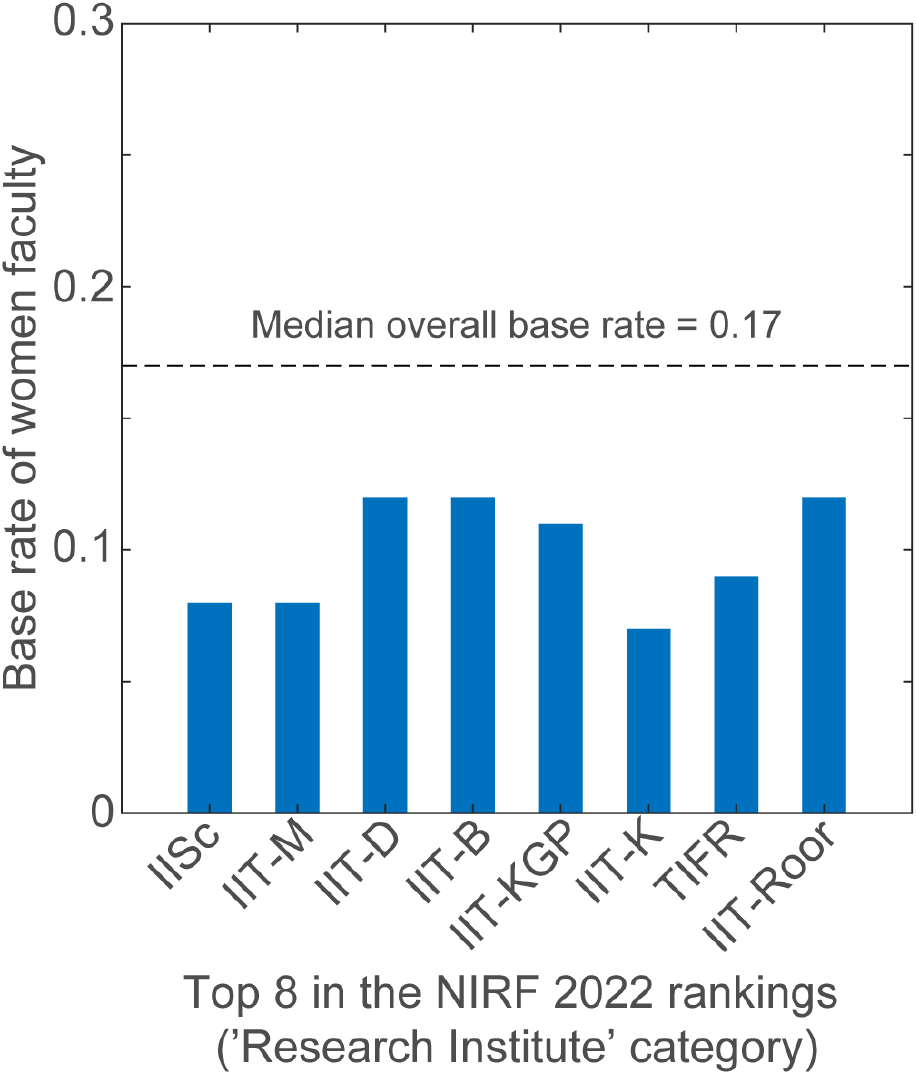
The base rates of top-ranked institutes/universities is below the median overall base rate. Bar plot of the base rate of the top 8 institutes in the ‘Research Institute’ category of the NIRF 2022 rankings. IISc - Indian Institute of Science; IIT - Indian Institute of Technology, M-Madras, D-Delhi, B-Bombay, KGP-Kharagpur, K-Kanpur, Roor-Roorkee; TIFR - Tata Institute for Fundamental Research.

Additionally, the proportion of women in academia has been historically low with measures to increase the number of women only recently introduced. We tested if this was the case in our dataset by documenting the career stage of the women faculty members surveyed. We observed that 46.3% of the women faculty members in the 45 institutes surveyed were early-career, 27.5% were mid-career and the remaining 26.2% were senior-career (**Fig. 3**).

**Figure 3.**
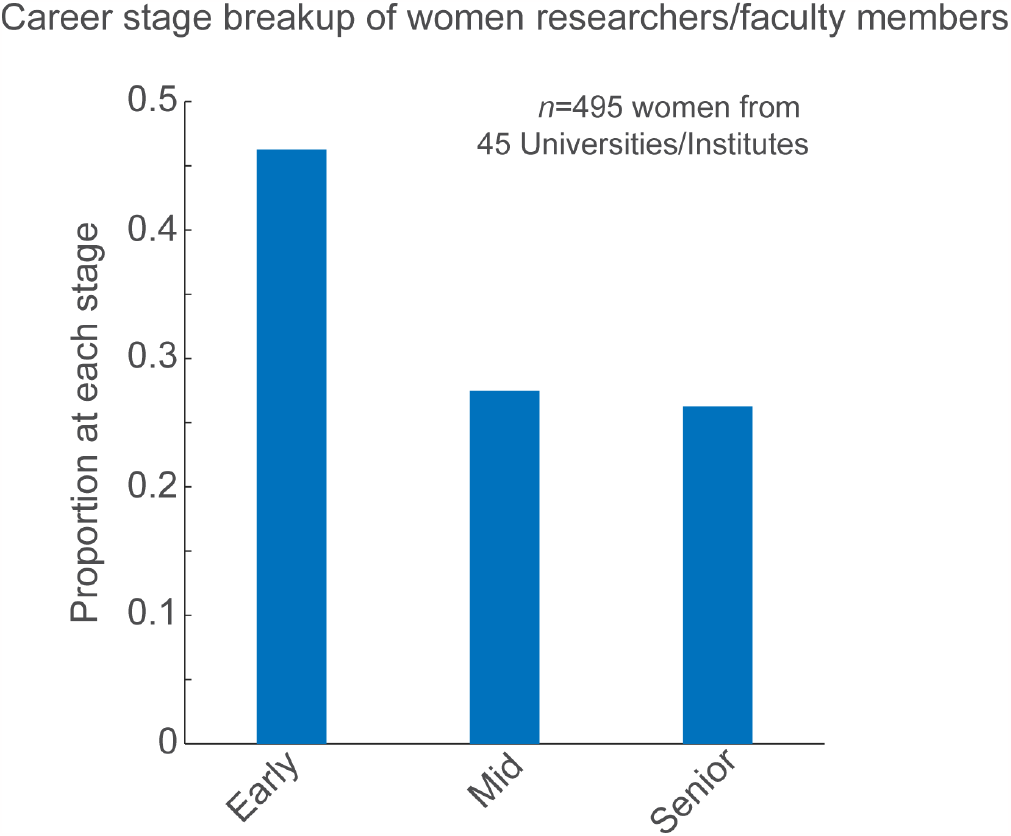
Proportion of women scientists by career stage. Of the 45 institutes surveyed and 495 women faculty members identified, 46.3% (229/495) were early-career, 27.5% (136/495) were mid-career, and 26.3% were senior-career.

### 2. Proportion of women speakers at women STEM conferences

Concomitantly, we sampled a total of 417 conferences/meetings for their women speaker ratios in two phases - Phase I: June 2020 to August 2021 (293 conferences) and Phase 2: August 2021 to March 2023 (124 conferences). All these events were either announced on Twitter or passed on to us by our Twitter followers. The median proportion of women speakers by field for conferences held in Phase I is plotted in **Fig. 4A**. We compared the proportion of women speakers in a conference against the base rate of the specific field to calculate the proportion of meetings with zero women speakers and meetings where the number of women speakers was below the base rate of women faculty in that field (**Fig. 4B**).

**Figure 4.**
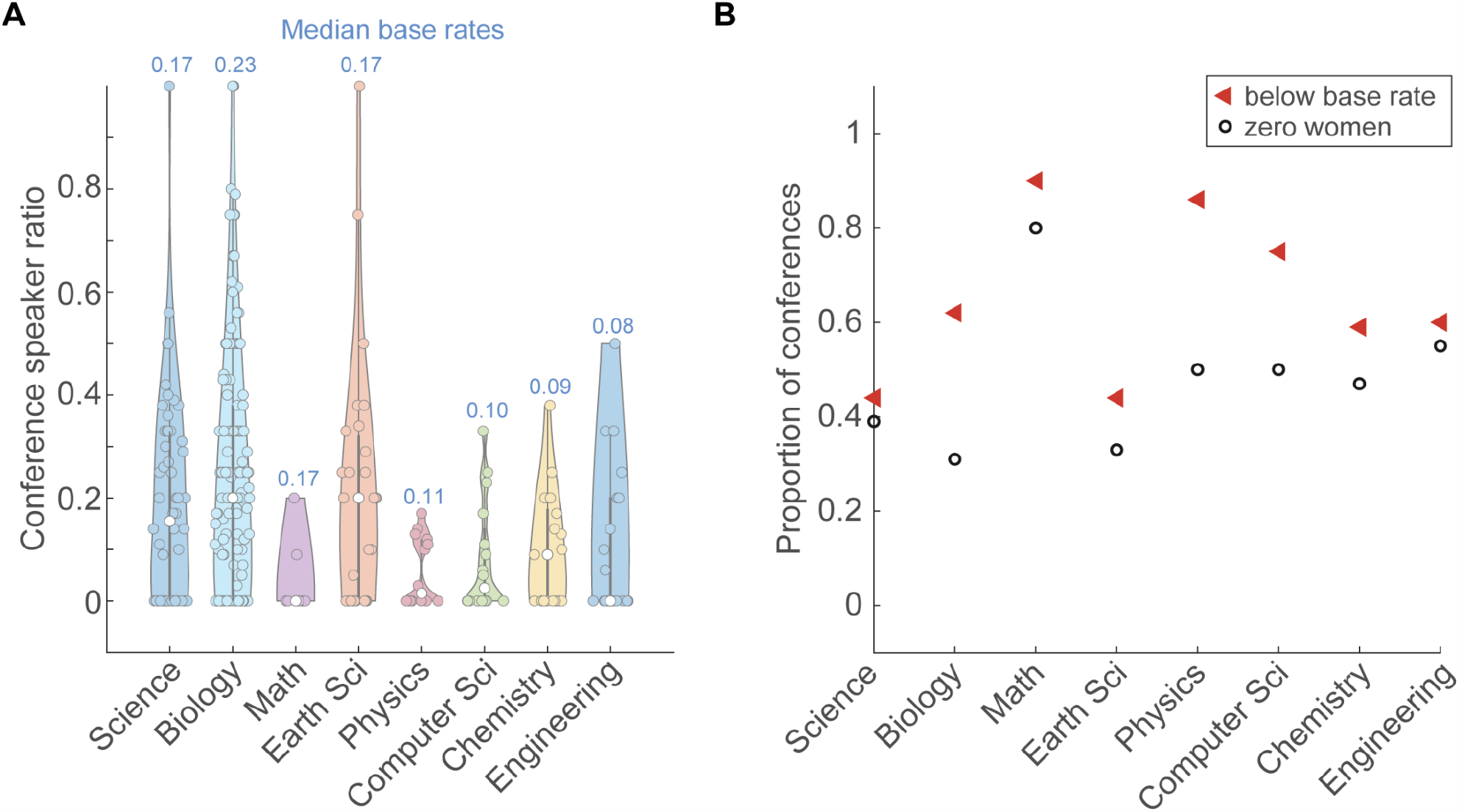
Phase I: Women are underrepresented in Indian STEM conferences. (A) Violin plot of the proportion of women speakers at Indian STEM conferences held between June 2020 and August 2021, organised by field. The median proportion of women speakers at these conferences is indicated by the white circle, and the median base rate for the corresponding field is mentioned above the plot. The thick gray lines represent the interquartile ranges. A total of 293 conferences were documented during this period, with n=54,137,10,27,14,16, 17 and 20 respectively for Science, Biology, Math, Earth Sciences, Physics, Computer Science, Chemistry and Engineering conferences. (B) Plot of proportion of conferences that featured zero women speakers (black circle) and the proportion of conferences that had women speaker ratios less than the base rate for the field (red triangle). For general Science conferences, Biology, Mathematics, Earth Science, Physics, Computer Science, Chemistry and Engineering, proportion of conferences with zero women speakers: 39%, 31%, 80%, 33%, 50%, 50%, 47% and 55%; proportion of conferences that underrepresented women compared to the base rate: 54%, 53%, 90%, 44%, 64%, 69%, 47% and 60%.

In Phase I, 80% of the conferences in Mathematics featured no women speakers, and overall 39% of all conferences conducted in this period had no women speakers. Similarly, 90% of Mathematics conferences featured fewer women than would be warranted by the base rate, and 60% of all conferences in Phase I underrepresented women.

In Phase II between August 2021 and March 2023, we documented 124 conferences, and the median proportion of women speakers by field for conferences held in this phase is plotted in **Fig. 5A**. Again, we calculated the proportion of conferences that had zero women speakers, as well as the proportion of conferences that underrepresented women (**Fig. 5B**). Like Phase I, Phase II had a significant number of conferences that underrepresented women, with 26% of all conferences in this period having no women speakers, and 55% of conferences with women speaker ratios below base rates for the respective field.

**Figure 5.**
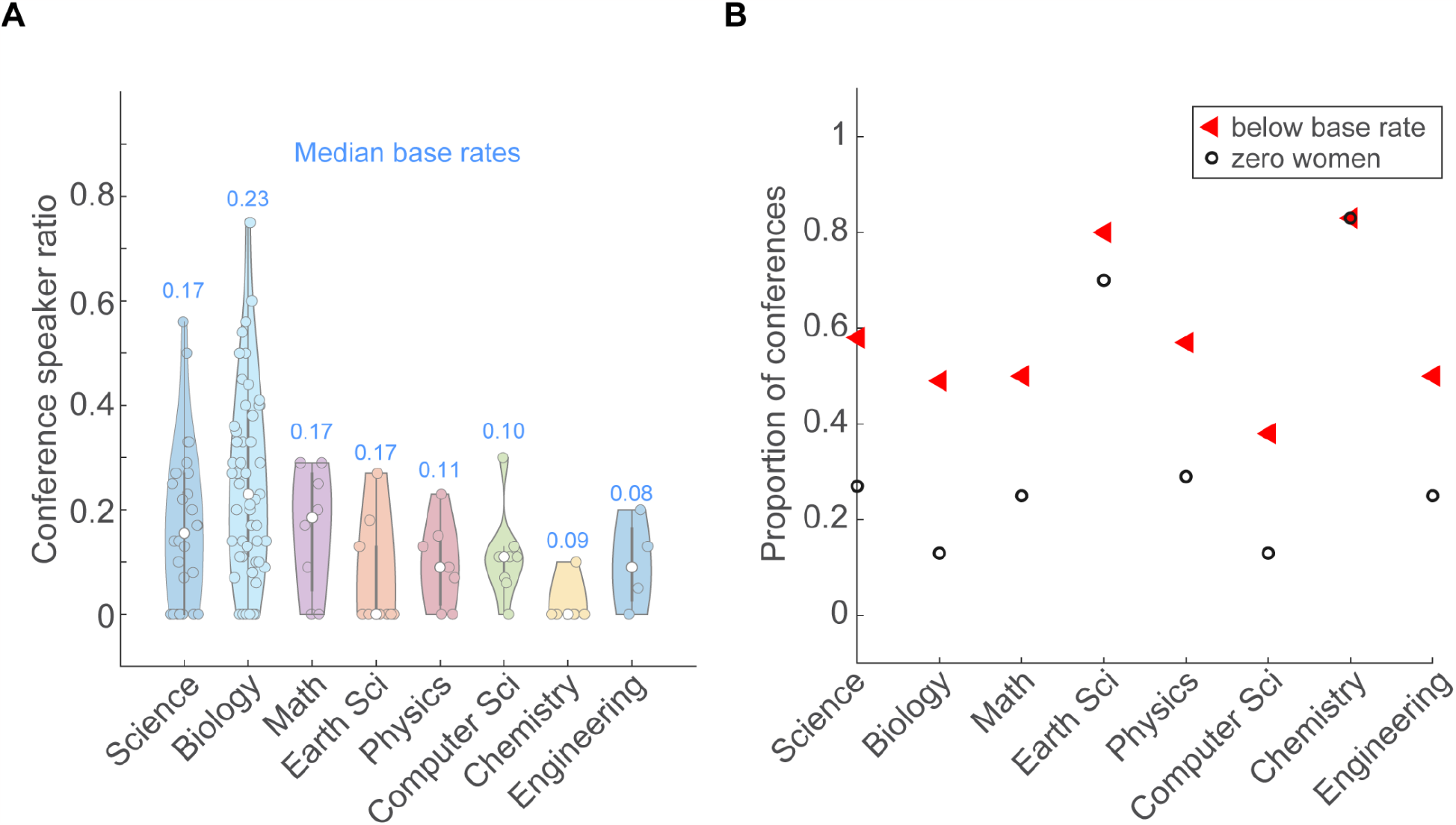
Phase II: Women continue to be underrepresented in Indian STEM conferences. (A) Violin plot of the proportion of women speakers at Indian STEM conferences held between August 2021 and March 2023, organised by field. The median proportion of women speakers at these conferences is indicated by the white circle, and the median base rate for the corresponding field is mentioned above the plot. The thick gray lines represent the interquartile ranges. A total of 124 conferences were documented during this period, with n=26,55,8,10,7,8,6 and 4 respectively for Science, Biology, Math, Earth Sciences, Physics, Computer Science, Chemistry and Engineering conferences. (B) Plot of proportion of conferences that featured zero women speakers (black circle) and the proportion of conferences that had women speaker ratios less than the base rate for the field (red triangle). For general Science conferences, Biology, Mathematics, Earth Science, Physics, Computer Science, Chemistry and Engineering, proportion of conferences with zero women speakers: 27%, 13%, 25%, 70%, 29%, 13%, 83% and 25%; proportion of conferences that underrepresented women compared to the base rate: 58%, 49%, 50%, 80%, 57%, 38%, 83% and 50%.

Comparing the women representation in Phase I and Phase II, we observed that Mathematics and Computer Science conferences fared better in Phase II with a reduction in the number of conferences that featured zero women speakers (80% and 50% in Phase I vs. 25% and 13% in Phase II respectively; **Fig. 6A**) and those that underrepresented women (90% and 69% in Phase I vs. 50% and 38% in Phase II respectively; **Fig. 6B**). However, other fields had an upturn in women speaker underrepresentation, such as Chemistry and Earth Science. It needs to be noted that the number of conferences that were documented for each field in Phase II was significantly lower in number and therefore statistical significance cannot be drawn based on this data.

**Figure 6.**
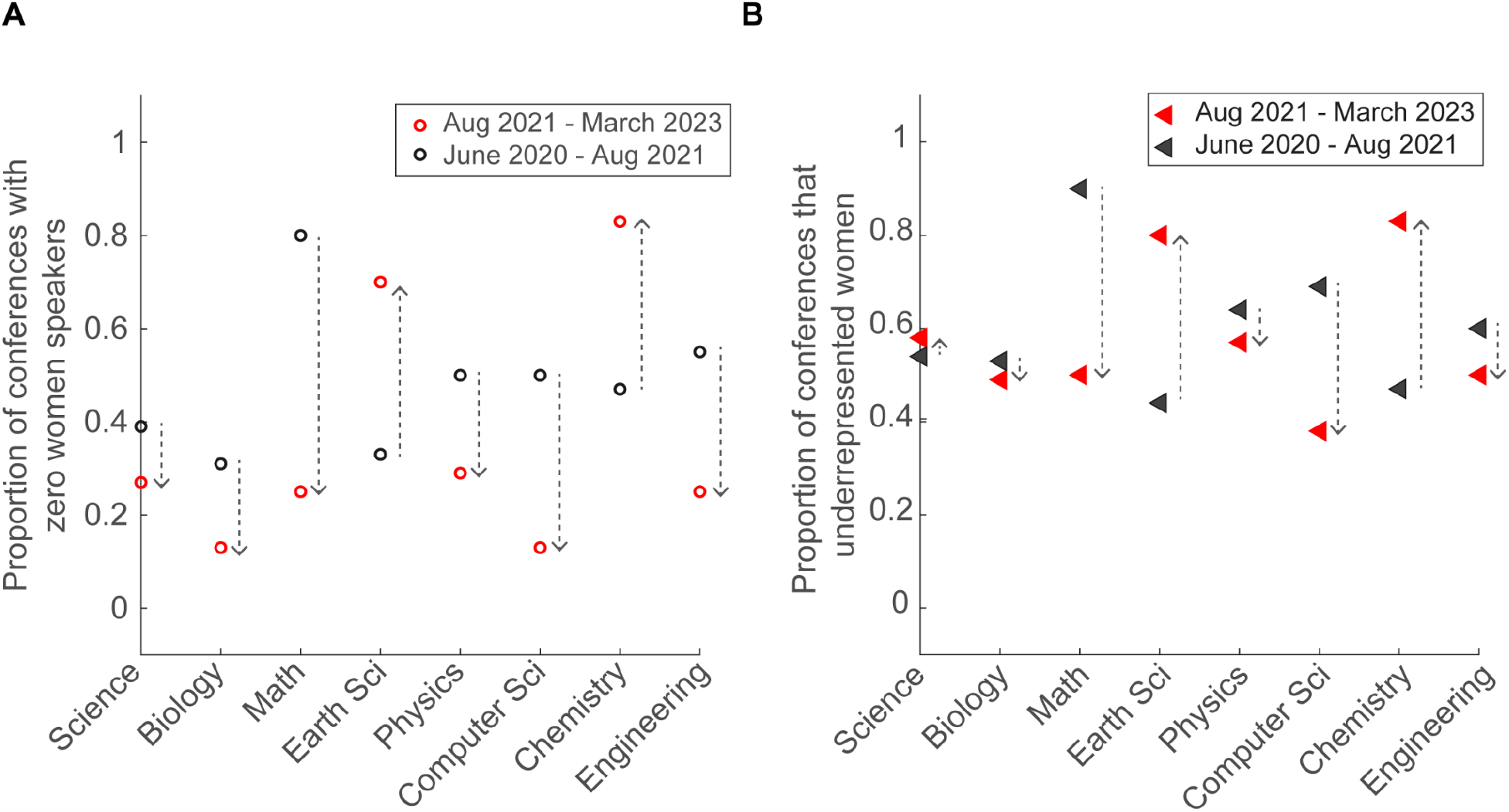
Comparison between Phase I and II for underrepresentation of women in Indian STEM conferences. (A) Plot of proportion of conferences that featured zero women speakers in Phase I (black circle) Phase II (red circle) (B) Plot of proportion of conferences in Phase I (black triangle) and Phase II (red triangle) that had women speaker ratios less than the base rate for the field. The dashed arrows show the increase or decrease in Phase II compared to Phase I.

Taken together, the data indicates that publicly tagging and calling out organizers and other invited speakers for low or zero women speaker representation seems to have a positive overall effect in the proportion of conferences with zero women speakers (46% in Phase I vs. 25% in Phase II). However, there is continued underrepresentation of women in conferences across all fields.

## Discussion

As of the time of writing this manuscript, we have not yet found a central Indian database that calculates and maintains subfield specific base rates of representation for Indian women faculty in Indian STEM academia. We found that NIRF reports are one way of collecting publicly reported data on women faculty representation. However, the obfuscation, incompleteness, and lack of inclusion of specific STEM fields in research, makes the data unusable to get a global view of changing trends in representation.

From our data, the differences we observe within the subfields may inherently reflect fields that are deemed socially acceptable for women in science (**Fig 1**). For example, Biology has the highest proportion of women faculty among all the subfields and Biology is traditionally considered a ‘soft science’^4^. Whereas Engineering, Chemistry, Computer Science and Physics show base rates of women faculty that hover around 0.1 - the lowest among all subfields. Even within prestigious institutions with high NIRF rankings such as IISc and multiple IITs (**Fig 2**), the proportion of women faculty remains unacceptably low. Moreover, the drastic decrease in the proportion of women faculty through the career stages (**Fig 3**) shows unmistakable attrition - also known as the ‘leaky pipeline’. Similar to what has been shown by other reports^5^, women faculty face multiple insurmountable barriers during their career progression that results in them quitting STEM academia for other careers and ventures.

We divided our data collection and analysis into two phases - Phase I from June 2020 to August 2021 (14 months **Fig 4**) and Phase II from August 2021 to March 2023 (8 months **Fig 5**). We calculated the proportion of conferences with lower than base rate along with events with zero women representation. Comparing Phases I and II (**Fig 6**), we found a decrease in the proportion of conferences with less than base rate representation of women speakers in all subfields except Earth Science and Chemistry where we found an opposite trend. Similarly, we observed a decrease in the proportion of conferences with zero representation of women speakers in all subfields except Earth Science and Chemistry.

Looking at other sources for similar data, we found that the UNESCO Institute for Statistics^6^ cites the percentage of Indian women researchers in science in 2015 at 13.9%. More recently, work by Swarup and Dey (2020)^7^ profiled the representation of women faculty in the top 20 Indian STEM institutes (ranked by NIRF) and found the average percentage to be 11.24%. Comparing these numbers to our calculated median rate of 16.6%, we see that the state of women faculty representation has not significantly changed at least over the last decade.

When we began BWI in June 2020, the world was in the middle of the COVID-19 pandemic. The resulting increase in virtual meetings and online presence made it easier to collect data on events, panels and discussions happening in Indian STEM academia. Twitter, in particular, was a useful tool not only to collect information and profile events, but also to interact with allies and gain access to events that were announced on social media platforms other than Twitter via reports by allies. More recently, with the changing goals and rules of X/Twitter^8^ as a social media platform, and with the gradual transition of events from being all-virtual to hybrid or all in-person, it is becoming difficult to maintain a stream of events that can be profiled.

Private messages or ‘Direct Messages’ (DMs) received from followers of the BWI Twitter handle show a wide variety of emotions. Most women and allies are thankful that a public-facing handle can rally against and call out systemic inequities along with providing base rate data for each subfield. Most women scientists across career levels are usually fearful of being vocal and visible in calling out systemic inequities. Especially, in Indian STEM academia, such an outspoken attitude costs women in terms of grants, collaborations, goodwill, and career-advancing steps such as promotions. In the last few years, BWI has acted as an anonymous repository for these women and allies to report Indian STEM events with minimal or zero women speakers. Comparatively, the few replies and DMs we receive from men have been dismissive of us and our efforts at best, combative at worst, even in public-facing replies and conversations.

A major learning from our experience of running BWI for 2 years is that data collection and dissemination is necessary to drive change. It is difficult to solve a problem as vast as discrimination without first defining a scope with the information available at hand. In 2021, there was no publicly available data source showing the proportion of women faculty in Indian STEM academia. We built this database and narrowed our scope to women faculty doing active scientific research in STEM institutes. Even though it was not possible to profile every single STEM institution across the country, we included a variety of institutions in our final list, such as - public and private universities, specialty biology and engineering institutes and interdisciplinary institutions such as Tata Institute for Fundamental Research. Profiling a total of 98 such institutes gave us an estimate of the proportion of women faculty members working in STEM academia across India.

As we used these base rates to call out events on Twitter, we found that a large proportion of our audience was not aware of low base rates of women in science and across subfields. Not only did this induce conversation around the topic, but also helped women and allies to notice and voice out concerns of under-representation in their own work environments.

One of our main continuing goals has been to think about how to extend similar data collection and analyses across other axes of bias such as other genders, caste, class and religion. While we have not been able to make significant progress on this front, we hope to better engage in the future and assist individuals/groups that take on this endeavour. A point of future work can be to collate data from the All India Survey on Higher Education (AISHE)^9^ so that a similar set of analyses can be extended to look at base rates and attrition among women PhD students and Postdoctoral Fellows. Overall, the transition from a Postdoc to a faculty member position (e.g., Assistant Professor) has been shown to be a major point of attrition for women in science^10^. This is because usually, this point in time coincides with family and social pressures that women face to leave the workforce and start families^11^. A more recent study indicates tenured senior women faculty are more likely to leave academia due to toxic workplace climates^12^.

Gender Advancement for Transforming Institutions (GATI) launched in February 2020 by the Department of Science and Technology (DST) is a pilot program that aims to document qualitative and quantitative gender data at the institutional level^13^. For example, <u>here</u>^14^ is a GATI report submitted by IIT-Delhi in January 2023. Unfortunately, since the program is in its nascent stages and because there is no external pressure to document and improve, we are yet to see the long-term effects of both data collection and eventual implementation at the policy level.

Taken together, we have observed and quantified that data collection and dissemination is necessary to drive change towards gender parity in Indian STEM academia. Given the difficulties in accessing gender data, we strongly believe that universities and institutes should perform regular gender audits and make the data easily and publicly available. Annual audit data should be tracked to follow trends and set goals. Our hope is that this effort eventually becomes self-driven and self-sustaining without external motivation from top-down orders from the Ministries or the Government.

As we (and others) have recommended in the past^15^ (and summarised below), increasing the proportion of women and minorities in Indian science requires clearly earmarked resources, and importantly, the will and strong commitment to equity from the leadership of Indian science agencies and universities/institutions.

### A. Support for postgraduates and advanced degree holders

1. <u>Abolish ageism</u> – Most early-career grants/positions require candidates to be below 35/40. This limitation needlessly penalises any career paths that do not follow default, traditional models originally built to facilitate men’s careers at the expense of women’s.
2. <u>Institute stable mentorships and support networks in each organisation</u> – Early-career women benefit from mentorship and more importantly, sponsorship from established academics in the field.

### B. Support for early and mid-career scientists

1. <u>Mandate the creation of an ‘Office for Equity and Inclusion’ in every institution</u> – These will act as hubs to connect all women in the institution and nucleate stable mentorships and support networks
2. <u>Provide support for families</u> – Provide adequate parental leave for both men and women. Account for childbirth in grant decisions and offer extensions when necessary.
3. <u>Institute tenure clock extension policies for women who need it due to childbirth</u> – This will ensure a level playing field for women scientists to reach important career milestones, such as tenure and promotion
4. <u>Ensure at least 30% women scientists are included on all panels</u> – Especially those that are related to career drives, recruitments, budget proposals and promotion to tenure. The active presence of women on such panels will provide much needed support to and understanding of decision-making.
5. <u>Set up a day-care centre on campus</u> – Childcare is not solely the mother’s responsibility. Providing childcare options on the academic campus can alleviate a major point of stress for new parents, especially women scientists who are new mothers.
6. <u>Be mindful of schedules of young parents (men and women)</u> – Do not exclude young parents or faculty members with family responsibilities by conducting official meetings beyond working hours.

### C. Support for senior scientists

1. <u>Promote and encourage long-term mentor-mentee relationships</u> – Early career women scientists should be supported with support, mentorship and sponsorship that can lead them through consistently successful careers. Long term, negative social biases must be mitigated with strong institutional policies.
2. <u>Prioritise including experienced women scientists’ voices in academy-, department- and government-level decisions</u> – Having women’s voices be heard will bring much needed diversity of thought and foster creative solutions to difficult problems
3. <u>Promote career-furthering activities such as sabbaticals</u> – Academic sabbaticals can help catalyse new directions and collaborations. Women academics generally hesitate to take sabbaticals due to societal pressures that require them to stay at home and take care of their families.
4. <u>Establish and regularly conduct gender sensitisation workshops for staff and faculty members at all levels</u> – Effectively challenging and changing generations’ worth of ingrained sexism is difficult. Therefore, regular training and workshops should be necessary for all staff and faculty members to initiate and maintain appropriate behaviour and attitudes.

## Acknowledgements

We would like to thank Leeba Ann Chacko, Mitali Shah, Harsh Kumar, Joel Joseph, Srivarsha Rajshekar, Srijani Biswas, Devashish Arvind Pande, Hradini Konthalapalli, Ravi Vishwakarma, Hymavathy Balasubramanian for data collection, Abhishek Chari, Sumeet Yamdagni for advice and discussions, members of our Advisory Board Ram Ramaswamy (IIT Delhi), Shobhana Narasimhan (JNCASR), Bittu Rajaraman (Ashoka University), Vidita Vaidya (TIFR Mumbai), Sandhya Visweswariah (SSV, IISc); SSV and Sandhya Koushika (TIFR Mumbai) for comments and suggestions on this manuscript; European Molecular Biology Organisation for funding; Ravinder Kaur (IIT Delhi) for the GATI self-assessment application; all our well-wishers who have consistently reported events that we missed.

